# Cerebral hemodynamics and cytochrome c oxidase in hypoxic-ischemic rabbit fetuses

**DOI:** 10.64898/2026.10.05.755935

**Authors:** Lorenzo Cortese, Giuseppe Lo Presti, Sergio Berdún, Gemma Bale, Zuzana Kovacsova, Elisenda Eixarch, Eduard Gratacos, Ilias Tachtsidis, Turgut Durduran

## Abstract

**Background:** Understanding fetal brain responses to hypoxia-ischemia is of growing interest, as it remains unclear to what extent these responses differ from those of newborns and whether developmental adaptations during gestation contribute to such differences.

**Materials and methods:** Using a combination of broadband near-infrared spectroscopy (bNIRS) and diffuse correlation spectroscopy (DCS), we measured brain microvascular oxygen saturation (StO_2_), total hemoglobin concentration (THC), relative cerebral blood flow (rCBF), and the oxidation state of cytochrome c oxidase (oxCCO) during an acute ischemic challenge in the brains of five rabbit fetuses in utero. Simultaneous measurements were acquired from four control fetuses located in the contralateral uterine horn.

**Results:** During the hypoxia-ischemia phase, we observed a rapid reduction of CBF (47% ± 29%) and StO_2_, (from 64.7% ± 1.5% at baseline to 40.6% ± 2.0%) together with an increase of oxCCO (3.7μM ± 0.5μM).

**Conclusion:** The simultaneous increase in oxCCO and decrease in rCBF and StO_2_ during hypoxia-ischemia reveals a cerebral response pattern that differs from that commonly reported in the newborn brain. These findings support the hypothesis that unique developmental adaptations influence fetal cerebral metabolism during hypoxic-ischemic stress.

## 1 Introduction

Hypoxic-ischemic encephalopathy (HIE) remains one of the leading causes of long-term neurological disability and death in the perinatal period [1], and substantial research effort continues to be directed at understanding the mechanisms of hypoxic-ischemic brain injury in the newborn [2,3]. A large proportion of infants who suffer HIE still go on to develop cerebral palsy, cognitive impairment, or epilepsy, underlining the continued need for a better understanding of how the newborn brain responds to oxygen and perfusion deprivation. It is now well recognized that a substantial proportion of hypoxic-ischemic brain injury diagnosed at birth in fact originates before labor, during fetal life, and this has driven growing interest in characterizing hypoxic-ischemic injury in the fetus rather than exclusively in the newborn [4].

While the metabolic response of the brain to hypoxia-ischemia has been extensively characterized in newborn animal models, including studies using broadband near-infrared spectroscopy (bNIRS) to continuously track hemodynamic and cytochrome c oxidase (oxCCO) changes as a direct marker of mitochondrial oxidative metabolism [5-8], comparatively little is known about how cerebral metabolism behaves in the fetus during an equivalent ischemic insult.

Most of the physiological data available on the fetal response to hypoxia-ischemia have come from invasive or indirect measures, such as blood gas sampling, microdialysis [9], or are based only on cerebral oxygenation [10]. Only few seminal studies on fetal sheep models [4,11,12] focused on continuously monitoring the cerebral hemodynamic and metabolic response to hypoxia-ischemia, with the ultimate objective of establishing how the timing, severity, and pattern of antenatal hypoxic-ischemic insults shape the resulting pattern of brain injury. These studies have highlighted significant differences of the fetal brain response respect to the newborn brain, involving adaptive mechanisms, such as redistribution of blood flow towards the brain and suppression of metabolic demand, that are specific to the fetal condition and that evolve over the course of gestation [12].

The aim of this study was therefore to use a diffuse correlation spectroscopy (DCS) system [13] together with a bNIRS system [14], combining continuous, non-invasive monitoring of cerebral blood flow with concurrent measurement of cerebral oxygenation and oxCCO [5,7,15], to investigate the hemodynamic and metabolic response of the fetal brain to a hypoxic-ischemic challenge in a rabbit model of fetal hypoxia-ischemia [16].

## 2 Methods

### 2.1 Experimental set-up and data analysis

A hybrid diffuse optical system consisting of a DCS module and a bNIRS module, was utilized. The DCS module consisted of a fiber coupled narrowband continuous wave laser at 785 nm (iBeam Smart, Toptica, Germany) as source, eight fiber coupled single photon counting avalanche photodiodes (SPCM-AQ4C, Excelitas, Canada) for detection (only two channels have been used, one dedicated to the case fetus, one to the control fetus), and an eight-channel autocorrelator board (Flex05oem8ch, Correlator Inc., USA) for reconstructing the intensity autocorrelation curves *g*_2_. In order to acquire simultaneously DCS from ischemic and control fetuses, the light source was divided in two branches of equal intensity by the mean of a fiber splitter. The cerebral blood flow index CBFi was retrieved by fitting the autocorrelation curves with the solution of the correlation diffusion equation for a semi-infinite medium and the relative cerebral blood flow rCBF was then obtained by normalizing the CBFi to its baseline value [13].

The bNIRS module consisted of a fiber coupled halogen lamp as white source (HL-2000-FHSA, Ocean Optics, USA) and a fiber coupled spectrometer (QE-Pro, Ocean Optics, USA) optimized for wavelengths between 600 and 1000 nm. Changes (Δ) in the concentrations of oxyhemoglobin (HbO2), deoxyhemoglobin (Hb), total hemoglobin (THC) and oxCCO were obtained by using the UCL algorithm described in [5] based on the modified Beer-Lambert law. Tissue blood oxygen saturation (StO_2_) was then obtained by using the second differential of the water absorption spectrum method [17].

A microcontroller gated the switching between DCS acquisitions (integration time 1 s) and bNIRS acquisitions (integration time 400 ms); to avoid crosstalk between the two technologies, the temporal resolution to complete the DCS-bNIRS acquisition cycle was set to 6.5 s (see also section 2.2).

DCS and bNIRS fibers were placed over the fetus head by means of a miniaturized 3D printed probe allocating the source and detector fiber bundles, with source-detector fiber tips separation of 7 mm. The source fiber bundle was composed of two multimode fibers (one for DCS, one for bNIRS) with diameter of 200 µm and a ferrule termination with diameter of 2.5 mm; the detector bundle consisted of one single mode fiber for DCS, with core diameter of 4 µm, and one multimode fiber for bNIRS, with core diameter of 200 µm, in a single ferrule termination with diameter of 2.5 mm. An identical 3D printed probe, but allocating only DCS fibers, have been used to monitor the rCBF of the control fetuses (i.e., bNIRS only measured the case fetus while the DCS had one channel over the case fetus and one channel over the control fetus).

### 2.2 Animal model and measurement protocol

Four New Zealand White pregnant rabbits at 30th day of gestation (term 31-32 days) were included in this study (Granja San Bernardo, Navarra, Spain). Animal housing, handling and all experimental procedures were performed in accordance with applicable regulation and guidelines and with the approval of the Animal Experimental Ethics Committee of the Universitat de Barcelona (ref. 04/17) and the competent authority Generalitat de Catalunya (ref. 9318).

A simple sketch of the rabbit hypoxia-ischemia model [16] and protocol implemented is represented in Figure 1 (a), together with a photo taken during the surgery (b). Two to three fetuses per dam were used. Fetuses were randomly assigned to hypoxia-ischemia (case fetuses, N=5) or control group (N=4). Animals were administered with subcutaneous buprenorphine 30 min before surgery (0.03 mg/kg and thereafter, medicated with ketamine and xylazine (35 mg/kg and 5 mg/kg respectively). Induction was performed by intravenous administration of propofol (5 mg/kg). An intravenous infusion of 2mg/ml ketamine and 0.6 mg/ml xylazine was used for maintenance of anesthesia (70-80 ml/h). Dams were oxygenated during the whole procedure (3 L/min oxygen, facemask). Body core temperature was maintained with an electric pad. Intramuscular progesterone (4.5 mg/kg) was administered for tocolysis prophylaxis. Heart rate, respiratory rate, oxygen saturation, body core temperature and reflexes of the dams were monitored through all the protocol duration.

**Figure 1.**
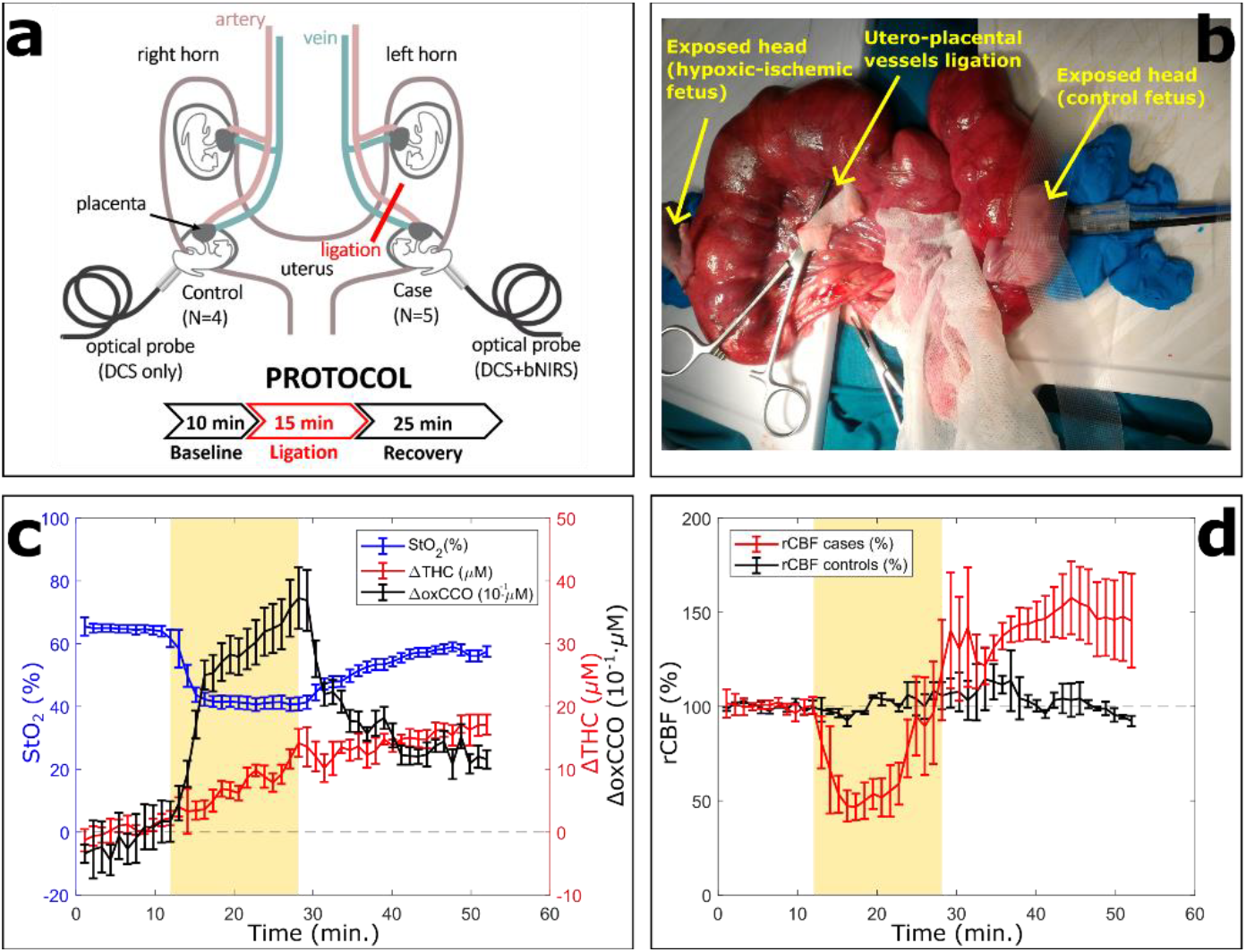
(a) Schematic representation of the uterus and of the protocol implementation; (b) photo the experiment showing two fetuses in two opposite uterine horns, with of the optical probe placed on the “exposed” head of a fetus and fixed with plasteline and surgical tape; (c) cerebral StO2, THC and oxCCO changes during the protocol; (d) rCBF of fetuses in the hypoxia-ischemia group (“cases”) and control group. Solid lines represent the average value among five fetuses (four fetuses in the case of the control group), error bars the standard error of the mean. Hypoxic-ischemic phase is highlighted in yellow.

The gravid uterus was exteriorized through a lower mid-line laparotomy. Uterine horns (one horn dedicated to ischemic fetus and the opposite to control fetus) were carefully manipulated in order to localize the orientation of the fetus inside the gestational sacs. A purse string suture was performed in the uterine wall and the fetal head was exteriorized through a 5 mm-incision hysterotomy, and fetal membranes were carefully dissected out.

The optical probes (one for the hypoxic-ischemic fetus, one for the control) were then placed onto the fetus head, in the parietal area 1-2 mm aside from the midline and were fixed with modelling clay and surgical tape (see Figure 1 top right). The duration of the complete measurement protocol lasted 50 minutes and was divided in three phases: a basal acquisition of ten minutes, the following fifteen minutes characterized of hypoxic-ischemic challenge, and finally the acquisition of twenty-five minutes posterior to the release of the ligation. The hypoxic-ischemic challenge in the fetuses of the ischemia group was obtained by clamping of utero-placental vessels of their own gestational sac. For this operation, vessels were carefully dissected out and clamped using a Kelly hemostat (13018-14, FST) for 15 min. Sterile gauzes soaked in ringer lactate solution covered the clamping surface to avoid damage of the vessels. After this period the clamp was released. Vessels of fetuses in control group were kept intact over all the duration of the protocol. Both uterine horns were continuously rinsed with sterile and warm ringer lactate solution. After the experimental protocol, pregnant rabbits were euthanized by means of pentobarbital (intravenous administration, 200mg/kg) and fetuses by decapitation.

We comment here that, due to the complexity of the surgical operation, for each fetus the vessel ligation required different maneuver times ranging from ten to forty seconds. In light of this, data have been down sampled to a temporal resolution of 65 s from the original resolution of 6.5 s by averaging 10 subsequent acquisitions, to compare and average results from different fetuses.

## 3 Results

Figure 1 (c) shows the microvascular cerebral StO_2_, ΔTHC and ΔoxCCO of the fetuses of the hypoxia-ischemia group during all the three phases of the hypoxia-ischemia protocol (baseline, ligation, recovery). Figure 1 (d) shows the rCBF of hypoxic-ischemic fetuses compared to control fetuses. Data reported are the mean values over five fetuses (four in case of the control group), and errorbars represent the standard error of the mean. Here, we would like to mention that this was a proof-of-principle exploratory study; given the small number of fetuses measured, the results presented are not statistically robust, and some features of the averaged blood-flow response may be influenced by individual animals and potential variability/movement artifacts in the completeness of ligation.

The ligation of utero-placental vessels caused a rapid decrease of CBF (47% + 29% with respect to basal condition) and StO_2_ (from 64.7% + 1.5% at basal condition to 40.6% + 2.0% at the end of the insult), while oxCCO increased (3.7μM ± 0.5μM above baseline value). After the ligation was releases, the monitored parameters started recovering towards the baseline values. The mean values of rCBF, StO_2_ and ΔoxCCO at the end of the recovery phase (twenty-five minutes after the ligation release) were respectively 143% + 13%, 57.5% + 1.5% and 1.3μM ± 0.2μM.

Figure 1 additionally shows an increase of rCBF before the release of ligation and the increase of THC, both during ligation and release phases, reaching a maximum increase of 17.1μM ± 1.6μM above baseline value at the end of the protocol.

Lastly, in the control fetuses (Fig. 1 (d)), no significant changes in CBF were detected during all the different phases of the protocol with respect to the baseline.

### 4 Discussion

As expected, the utero-placental vessel occlusion abruptly reduced the fetus’s supply of oxygenated blood (hypoxic-ischemic model), causing a rapid reduction of cerebral blood flow and cerebral blood oxygen saturation. Differently from case-fetuses, brain hemodynamics of fetuses of the control group (only rCBF monitored) was not affected by the protocol.

In hypoxic-ischemic fetuses, we also noted an increase of oxCCO during the hypoxic-ischemic insult. This result agrees with the results of A. Gunn [4,11,12], which reported an oxidation of cytochrome c oxidase in the brain during hypoxia-ischemia in fetal sheep generated by umbilical cord occlusion. The increase of oxCCO during the ischemia reflects a reduction/depletion in substrate delivery which reduces the transfer of reducing equivalents from the tricarboxylic acid cycle, suggesting a suppression of the metabolic activity. This could be interpreted as an active, regulated adaptation of the brain to low oxygen levels, downregulating the metabolic demand to avoid permanent damage (“adaptive hypometabolism”), or as biochemical malfunction, being the electron chain unable to donate electrons effectively during the ischemia.

These results strongly differentiate the fetal brain behavior from newborn. In newborn animal models of hypoxia-ischemia (almost all piglet-based), the oxCCO response is typically a decrease during hypoxia-ischemia challenges, reflecting impaired oxidative phosphorylation and failing mitochondrial function [4-8,18]. When oxygen tension at the mitochondria falls to a critical level there is a coincidental reduction of the electron transport chain and fall in the phosphorylation potential, assuming that there are electrons feeding the chain, and consequently cytochrome c becomes almost fully reduced. On the contrary, suppression of metabolism on fetal brain (i.e., increase of oxCCO) can be interpreted as a protective adaptation that helps explain why the fetus can tolerate remarkably long periods of severe hypoxia-ischemia without the same degree of injury seen in more mature or newborn brains. Previous studies demonstrated that this protective adaptation of metabolism is also gestation-dependent: Drury et al. [12] showed that the rate of cytochrome oxidation during profound asphyxia becomes faster with advancing gestational age in fetal sheep, and interpreted the loss of this protective, slower/adaptive redox response as one mechanism contributing to the fetus’s increasing vulnerability to injury as pregnancy progresses.

Figure 1 additionally highlights two interesting trends: the recovery of CBF before the release of ligation and the increase of THC, both during ligation and release phases. This could suggest the presence of compensatory mechanism in the brain of the fetus in reaction to hypoxia-ischemia.

## 3 Conclusion

The combined measurement of cerebral blood flow, oxygenation and metabolism provides valuable insight into fetal hypoxia-ischemia. During the hypoxia-ischemia phase, we observed an increase of oxCCO together with a decrease of cerebral blood flow and oxygenation. These findings suggest that the biological and metabolic response to hypoxic-ischemic insults differs between newborn and fetuses. Differently from newborns, in fetuses we observed a protective mechanism regulating the adaptation of the brain to low oxygen levels by a suppression of metabolism. This distinction may be important for future fetal monitoring studies.

## Acknowledgement

This study was funded by: Fundación Cellex; FUNDACIÓ Privada MIR-PUIG; Horizon 2020 Framework Programme (688303, 871124, 101016087, 101017113, 101099291, 101099093); European Social Fund Plus; European Regional Development Fund (EFRE-OP 2014-2020 - Project no. 2021 FE 9032); Agencia Estatal de Investigación (PID2019-106481RB-C31/10.13039/501100011033, PID2023-151973OB-I00 PHOTOSHOCK, PID2023-147553OB-I00 SCOSWear, LUX4MED/MEDLUX, PHOTOMETABO); Fundación Carmen y Severo Ochoa (CEX2019-000910-S); Centres de Recerca de Catalunya; Agència de Gestió d’Ajuts Universitaris i de Recerca (2019-FIB-00968, 2022-SGR-01457); Institució Catalana de Recerca i Estudis Avançats (RIS3CAT-001-P-001682 CECH); Departament d’Empresa i Coneixement, Departament de Salut (SLT041/25/000082, SLT041/25/000083), Generalitat de Catalunya; National Institutes of Health (R21EB031261)

## Conflict of Interests or disclosures

Turgut Durduran is an inventor on relevant patents. Potential financial conflicts of interest and objectivity of research have been monitored by ICFO’s Knowledge & Technology Transfer Department. No financial conflicts of interest were identified. Ilias Tachtsidis is a Professor at University College London (UCL) and also the Founder and CEO of MetaboLight Ltd. The work presented in this publication was conducted entirely as part of his academic role at UCL. MetaboLight Ltd operates in a different field, and there are no financial or research conflicts of interest related to this work. The authors declare no additional conflicts of interest.

